# Frugivory and seed predation of fishtail palm (*Caryota mitis* Lour.) on the remote oceanic island of Narcondam, India

**DOI:** 10.1101/2023.04.06.535840

**Authors:** Abhishek Gopal, Sartaj Ghuman, Vivek Ramachandran, Navendu Page, Rohit Naniwadekar

## Abstract

Oceanic islands, due to their evolutionary history and isolation, hold a disproportionately high proportion of endemic species. However, their evolutionary history also makes them vulnerable to extinctions, with most known extinctions occurring on islands. Plant-animal interactions are particularly important on islands, as island systems generally have low redundancy and are more vulnerable to disruption either via extinction or by invasive species. Here, we examined the fruit removal and seed predation of a keystone palm, *Caryota mitis*, on the remote oceanic island of Narcondam. The island endemic Narcondam hornbill (*Rhyticeros narcondami*), was the sole seed disperser of the *Caryota mitis* (90 hours; N = 15 trees), indicating a lack of redundancy in seed dispersal of the palm on this island. While the invasive rodent, *Rattus* cf. *tiomanicus* was the sole predator of the *Caryota mitis* seeds in the forest (N = 15 individual fruiting palms, 416 trap nights). Overall, 17.1% of the seeds placed (N = 375 seeds) were removed. Seeds placed under and away from the canopy, and at different densities (2 plots with 10 seeds each; 1 plot with 5 seeds, respectively), showed similar removal rates. This is indicative of ambient seed predation and the lack of safe sites for the regeneration of *Caryota mitis*, with potential deleterious effects on the subsequent stages of the “seed dispersal cycle”. Here, from a data deficient site, we provide baseline information on the plant-frugivore interaction of a keystone palm and the potential impacts by an invasive rodent.

## 1. INTRODUCTION

Globally, most known species extinctions have occurred on islands (Bellard et al. 2016). Loss of endemic species and introduction of invasive species can have cascading impacts on key ecosystem processes, like seed dispersal and seed predation (Heinen et al. 2023; Moser et al. 2018). These critical ecosystem processes in tropical forests ensure plant regeneration and maintenance of plant diversity (Wang & Smith 2002). The initial stages of the seed dispersal cycle, namely, fruit removal and seed predation, determine the spatial template of the distribution of seeds in the forest (Wang & Smith 2002, Beckman & Rogers 2013). The disruption in these early stages of the dispersal cycle can have downstream consequences on the recruitment stage of the plants.

Islands have a smaller assemblage of frugivores compared to mainlands and thus endemic frugivores can play an irreplaceable role in seed dispersal. Similarly, introduction of invasive rodents (in absence of natural predators) can result in negative impacts on plant regeneration via seed predation (Auld et al. 2010; Meyer and Butaud, 2009; Traveset et al. 2008). Therefore, it is critical to determine the role of endemic frugivores in the seed dispersal process and the impacts of invasive rodents on seeds of native plants, especially those that are vital for endemic frugivores.

Palms (Family: Arecaceae) are a quintessential component of tropical wet forests, whose seeds are generally dispersed by animals (Couvreur & Baker 2013, Snow 1981). They are known to be negatively impacted by invasive rodents on islands (Auld et al. 2010, Meyer & Butaud 2009) to the extent that rodents have been responsible for the local extinction of palms on certain islands (Athens et al. 2002, Athens 2008). Notably, palms are an important fruit resource for frugivores on islands (Naniwadekar et al. 2021, Zona & Henderson 1989). Many palm species provide copious amounts of fruits and have staggered fruiting, providing frugivores ample fruit resources over a prolonged time period (Adler & Lambert 2008, Schaefer et al. 2002). This makes it interesting and important to determine seed dispersers of palms and the impacts of invasive rodents on palm seeds on oceanic islands.

In this study, we focus on the *Caryota mitis* Lour., a sympodial species of fishtail palm, to determine patterns of frugivory and seed predation on a remote oceanic island, Narcondam (area: 6.8 km^2^; elevation range: 0–710 m ASL; 13.4456°N, 94.2633°E) in the Andaman sea in south Asia. Insights from mainland studies show that *Caryota mitis* is consumed by over fourteen vertebrate species, both birds and mammals (Quek et al. 2020). It is widely distributed and abundant in south-east Asian tropical forests. It is hapaxanthic, wherein, post an initial vegetative phase, all its resources are exhausted in a prolonged flowering phase (Uhl and Dransfield 1987). They produce substantial infructescence in a basipetal sequence, thereby providing a continuous resource to frugivores and seed predators (Quek et al. 2020). One study estimates up to 805 fruits per infructescence (Quek et al. 2020).

On Narcondam Island, *Caryota mitis* is one of the most abundant plant species of more than 210 flowering plant species recorded from the island ( Table S1; Page et al. 2020). Like other palms, due to its staggered fruiting, it is likely to be an important resource for frugivores and seed predators on the island. This palm was found to be one of the most important large-seeded fruits in the diet of the point-endemic Narcondam Hornbill (Naniwadekar et al. 2021). The island is a relatively young volcanic island (~700,000 years old; Bandopadhyay, 2017) and is unique due to the absence of small-bodied frugivores like bulbuls (Pycnonotidae), barbets (Megalaimidae), and Asian Fairy Bluebird *Irena puella*. These are important seed dispersers in mainland Asia. The island, however, has relatively larger-bodied avian frugivores, including the point-endemic Narcondam Hornbill (*Rhyticeros narcondami*), two species of Imperial pigeons, *Ducula bicolor* and *Ducula aenea*, Asian Koel *Eudynamys scolopacea*, and Common Hill Myna *Gracula religiosa* (Naniwadekar et al. 2021). Frugivorous bats have also been reported from the island, which can play an important role in the long-distance dispersal of plant species (McConkey & Drake 2015). Unlike some other Pacific Ocean islands (Carpenter et al. 2020), there are no known extinctions of seed predators or dispersers from the island. Given the importance of this palm for the point-endemic hornbill, it is important to evaluate the potential impact of invasive rodents on the palm.

The Narcondam Hornbill is endemic to the small island of Narcondam and occurs in very high densities (~151 birds per km^2^; Naniwadekar et al. 2021). It is a central frugivore in the plant-frugivore community on the island (Naniwadekar et al. 2021). The oceanic island has a high abundance of rodents *Rattus* cf. *tiomanicus* (Ramachandran et al. unpublished data), which are likely to have been inadvertently introduced by humans. There have been a few previous expeditions to the islands which have reported a high abundance of invasive rodents, however, their ecological role has not been examined in detail (Shankar Raman et al. 2013, Vivek & Vijayan 2003). This provided an interesting setup to examine the role of plant-animal interactions and any potential role of invasive rodents. Here, using *Caryota mitis* as a focal taxon, we aimed to identify and determine the relative importance of different frugivores in its seed dispersal by conducting focal tree watches; and to determine the impact of the invasive rodent on the seed predation of the palm via an experimental framework wherein seeds were laid at different densities (high and low) and at different distances from the fruiting individuals (under the canopy and away from the canopy). We expected that we would not find Janzen-Connell (Connell 1971, Janzen 1970) effects due to the high abundance of invasive rodents on the island, resulting in ambient seed predation and a lack of safe sites for plant regeneration.

## 2. METHODS

To determine the main seed dispersers of *Caryota mitis*, we observed 15 individuals of fruiting *Caryota mitis* between 0600 hr to 1200 hr. During the fruit tree watches, we recorded frugivore species identity and the number of birds that visited the fruiting *Caryota mitis*. We also counted the number of fruits swallowed, pecked and dropped by the frugivores to estimate the fruit removal rate per visit. Additionally, we carried out night tree watches on 5 individual palms for 150 min over 3 nights to determine if any of the bat species on the island dispersed the seeds of *Caryota mitis*.

To estimate the seed removal rates by the invasive *Rattus* cf. *tiomanicus* and other animals, we set up four seed plots (1 m × 1 m), each under 15 fruiting *Caryota mitis* palms. We selected trees with relatively high fruit crop size and ensured the minimum distance between the selected palms was greater than 30 m. One exclosure plot and one seed plot were established below the parent fruiting tree along with two additional plots 15 m away in opposite directions of the parent tree. The plots below the parent tree (Exclosure and Below: high density) and one of the plots 15 m away from the fruiting tree (Away: high density) had ten seeds each. The other plot 15 m away (Away: low density) had five seeds. All the leaf litter was removed from within the plots. Under the parent tree, we did not alter the number of fruits/seeds but only marked and tagged 10 seeds in the plot for ease of monitoring. The number of seeds in away-high density plots was decided based on earlier studies (Gopal et al. 2021; Krishnan et al. 2022; Sidhu et al. 2015). For low-density plots, we have used the number of seeds that can be scatter-dispersed by hornbills in tropical forests (Naniwadekar et al. 2021). A free 50 cm Dacron fishing line was attached to the seed surface of each seed with a non-toxic super glue (Loctite® Super Glue Ultra Gel ™) to track seed fate following Gopal et al. (2021) and Sidhu et al. (2015) to ease detectability of marked seeds that are moved or predated. The seed plots were monitored weekly, and seed fates, intact (seeds with no visible signs of predation), predated (seeds with bite marks or with remains), and removed (seeds missing from within 10-m radius of the plot), were recorded following Gopal et al. (2021). Whenever seeds were missing from the plot, two observers (AG and SG) looked for the seeds in a radius of 10 m surrounding the plot. For further analysis, we merged the seed fates ‘removed’ and ‘predated’. Please see Table S3 for details of all the seed fates. The plots were monitored till all the seeds were removed or until the end of the study period. Camera traps were placed under the canopy of the 15 fruiting *Caryota mitis* trees (the same trees under which seed plots were established; Below: high density plot) to determine visitor diversity and visitation rates. The total camera trap effort across the 15 individual fruiting palms was 416 nights (range: 25–33; Table S2). We used a Generalised Linear Model with a binomial error structure to determine whether the proportion of seeds removed differed among the three categories (Below: high, Away: high and Away: low). Additionally, we opportunistically examined the seed predation of six other tree species fruiting during the study period (total N = 16; *Aphanamixis polystachya, Canarium euphyllum, Chionanthus sp., Codiocarpus andamanicus*, *Endocomia macrocoma*, and *Planchonella longipetiolata*). For these species, we laid out two seed plots, one under the canopy (Below: high density) and one away (15 m) from the canopy (Away: high-density) with 10 marked and tagged seeds in each plot. Camera traps were placed under the canopy plot. Please see Table S2 for additional details.

## 3. RESULTS

We observed Narcondam Hornbill in 11 of the 15 focal palm watches, while Asian Koel and Green Imperial-pigeons were observed only in one focal tree watch. The mean (± SE) visitation rate of Narcondam Hornbill on *Caryota mitis* was 0.23 (± 0.06) birds per hour. The Narcondam Hornbill was observed handling 49 fruits, of which 71.4% were swallowed, 18.4% were dropped, and 10.2% were inspected. The mean (± SE; range) fruit removal rate per visit was 3.5 (± 1.5; 0-16) fruits per visit. This includes only those fruits that the birds swallowed. During the focal palm watches, there were no observations of other frugivores handling fruits, including the Asian Koel and Green Imperial-pigeon that visited the focal palm trees. Additionally, in the night tree watches, we did not observe any frugivorous bats visiting *Caryota mitis*.

In 416 trap nights, the invasive *Rattus* cf. *tiomanicus* was found to be the sole vertebrate seed predator of the selected tree species on the island. The mean (± SE) visitation rate of the rodents was 1.75 (± 0.26) visits per trap night.

Despite thorough searches within 10 m of the plot, we did not find any evidence of caching or secondary seed dispersal. Of the 77 seeds of *Caryota mitis* that were removed, we either found predated seeds with fishing lines or fishing lines without seeds (N = 41), or we were unable to find the fishing line at all (N = 28).

For *Caryota mitis*, all the seeds in the exclosure plots were intact during the study period. However, for the other treatments, overall, 17.1% (SE = 3.06; range = 0–1; N = 375 seeds) of the seeds were removed. The proportion of seeds removed was similar across treatments, suggesting ambient predation (Table 1; Fig. 1). The other tree species had more than 50% of the seeds predated, and a very low overall removal rate of 8% was observed. More than 90% of the seeds of *Codiocarpus andamanicus* and more than 85% of *Planchonella longipetiolata* seeds were predated across treatments (Table S3). On the other hand, while more than 70% of the seeds of *Canarium euphyllum* were intact, 70% of the seeds of *Aphanamixis polystachya* were removed (Table S3). *Planchonella longipetiolata* was the only species for which we found evidence of invertebrate predation, with 13% of seeds (out of 100 seeds) being predated by invertebrate seed predators (5% and 8% in Below: high density and Away: high-density, respectively). However, there was no difference in the proportion of seeds that were removed between the plots under and away from the canopy, as inferred from overlapping SEs (Fig. S1).

**Figure 1.**
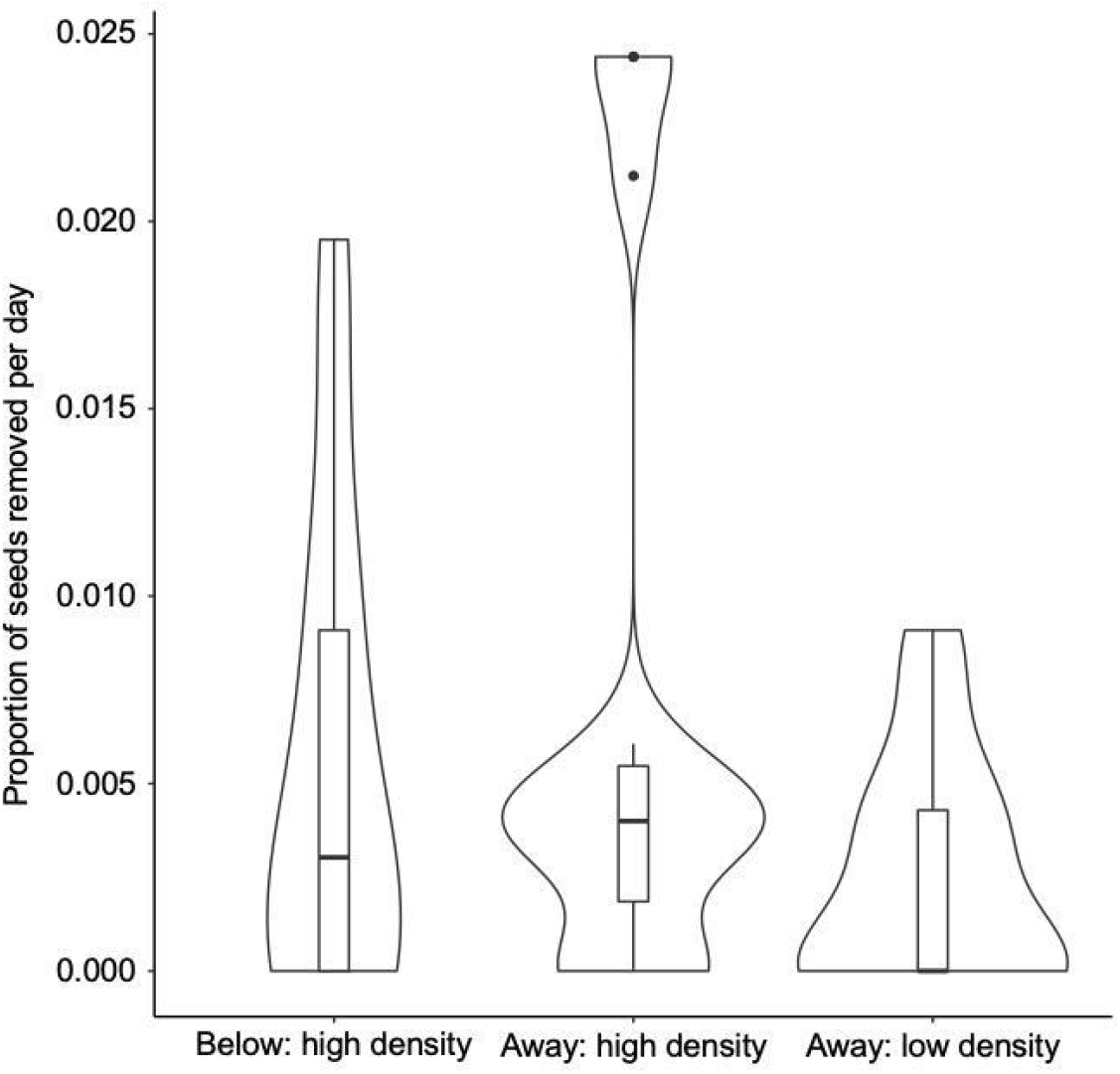
Violin plot (with box plots inside) depicting the proportion of seeds removed per day of *Caryota mitis* (*N* = 15) across the three categories - under the parent plant (Below: high density, 10 seeds per plot), away from the parent plant with high seed densities (Away: high density, 10 seeds per plot) and away from the parent plant with low seed densities (Away: low density, 5 seeds per plot).

**Table 1.** Summary of the generalised linear model with binomial error structure examining the relationship between the proportion of seeds removed across the three categories. The three categories are below the parent plant, away from the parent plant with high seed densities and away from the parent plant with low seed densities.

|  | Estimate | SE | *z* | *p* |
| --- | --- | --- | --- | --- |
| Intercept (Away: high density) | -4.961 | 3.107 | -1.597 | 0.11 |
| Away: low density | -1.060 | 6.105 | -0.174 | 0.862 |
| Below: high density | -0.257 | 4.701 | -0.055 | 0.956 |

## 4. DISCUSSION

Here, from a data-deficient region of the Indian Ocean, we provide baseline information on the frugivory and seed predation of a keystone palm species. In terms of frugivory, unlike the mainland sites (Quek et al. 2020), in 90 hours of observations, only the point-endemic Narcondam Hornbill was seen dispersing the seeds of the *Caryota mitis*. In terms of seed predation, the invasive rodents removed seeds in a pattern not consistent with the expectations of the JC hypothesis. Rodents removed seeds both near and away from the parent tree and in both low and high densities, indicating ambient seed predation by the invasive rodents on the island.

*Rattus* cf. *tiomanicus*, although a species native to south-east Asia, is known to be invasive in certain parts of its native geographic range and is also very abundant in the secondary forests and oil palm plantations in south-east Asia (Chinkok et al. 2007, Gibson et al. 2013). The photo capture rates of *Rattus* on Narcondam Island were 2.9 times higher than a mainland site in southern India, where seed plots were laid out in fragmented forests under four different fruiting tree species (Gopal et al. 2021), indicating a higher predation pressure on the island. Future research needs to examine the impacts of rodent removal on palm regeneration on the island.

We also sampled four other tree species, but unlike *Caryota mitis*, we could not find sufficient replicates for those tree species. However, our preliminary data for four other plant species also show similar seed removal rates near and away from the parent tree (Fig. S1), indicating the pervasive impacts of the invasive rodents as seed predators. Unlike *Caryota mitis*, the seed removal rates were much higher for these species. The variation in seed removal rates across the different plant species points towards variable seed predation pressure, which could potentially alter the recruitment trajectory of plants on the island in the long term. All the tree species we studied are important food plants of the point-endemic Narcondam Hornbill (Naniwadekar et al. 2021). It needs to be determined with more intensive sampling if the predation of seeds has cascading downstream effects.

Interestingly, we only found the Narcondam Hornbill feeding on the fruits of *Caryota mitis*. Frugivorous bats can also potentially disperse the seeds of *Caryota mitis*, however, we did not observe them feeding on the palm during our night tree watches. Imperial pigeons have been reported to disperse the palm seeds elsewhere (Quek et al. 2020), yet during our two-month stay on the island, we only found Imperial pigeons feeding on small-seeded plants (Naniwadekar et al. 2021). Among the large-seeded plants on the island, *Caryota mitis* had high species strength values highlighting its importance for the hornbill and in the community (Naniwadekar et al. 2021). Due to its relative abundance and long, staggered fruiting period, *Caryota mitis* is an important food resource for the hornbill (Naniwadekar et al. 2021).

There have been several previous expeditions to Narcondam that focused on the ecology of the point endemic hornbill. Although some do anecdotally report high densities of rodents (Shankar Raman et al. 2013, Vivek & Vijayan 2003), there have been no systematic efforts to estimate the densities of rodents and document their impact on the island biota. Although a before-after control-impact framework would have been ideal to conclusively examine the effects of the rodents, given the remoteness and ruggedness of the island, such information is lacking. Here we provide some baseline information on the potential impact of the invasive rodents on the islands. Invasive rodents have been shown to have deleterious impacts on other island biotas such as invertebrates, birds, and crustaceans (Harper and Bunbury, 2015). More intensive work is needed to systematically document the effect of these rodents on the island biota. Robust baseline data will help to identify the need for a management plan, in collaboration with the Forest Department and researchers, to control the invasive rodent populations on the island using established methods and systematically documenting the effects of rodent removal on vegetation recovery as has been done elsewhere (Miller-ter Kuile et al. 2021).

Invasive species have caused catastrophic declines and extinctions of several native and endemic species, especially on islands. Due to their isolation, small size and unique evolutionary history, island systems are more susceptible to such invasions. Narcondam Island, by virtue of its small size, remoteness and lack of terrestrial mammals, exemplifies these threats. Further studies are required to understand the population density and dynamics of *Rattus* cf. *tiomanicus*, and to resolve the biogeography and colonisation history of this species on the island. Systematic documentation and baseline information is needed on the densities of rodents and other key biotas on Narcondam Island to critically examine the impacts of the invasive rodent on the island biota and ecosystem of this unique island system.

## ACKNOWLEDGMENTS

We thank the Andaman and Nicobar Forest Department for giving us the permit (No:WII/NVP/NARCONDAM/2019). We thank D. M. Shukla (PCCF, Wildlife), A. K. Paul and Soundra Pandian for giving us the necessary permissions and facilitating our work. We thank Dependra Pathak, DGP (A&N), for giving us the required permissions. We thank Commandant A. K. Bhama and Captain Kundan Singh from the Indian Coast Guard for giving us permission and support. We thank Abhishek Dey, DC (South Andamans) for giving us permission. We thank Kulbhushansingh Suryawanshi, Divya Mudappa, T. R. Shankar Raman and G. S. Rawat for their support. We are thankful to the staff of the Special Armed Police unit led by Usha Rangnani (SP) for providing us with all the logistic help. We thank Elrika D’ Souza, Evan Nazareth, Rachana Rao, and Rohan Arthur for their support in Port Blair. AG thanks Hannah Krupa for helping with proofreading the manuscript.

## FINANCIAL SUPPORT

Funding and support were received from Wildlife Conservation Trust, IDEAWILD, Nature Conservation Foundation, Mr. Uday Kumar, M. M. Muthiah Research Foundation, Mr. Rohit and Deepa Sobti, and Mr. Aravind Datar.

## COMPETING INTERESTS

Authors have no competing interests to declare.

## ETHICAL APPROVAL

The study was observational and did not involve handling of the animals. The ethical clearance for the study was obtained from the Nature Conservation Foundation (NCF-EC-16/11/19-(43)). Additionally, we obtained the necessary permissions from the Forest Department (No: WII/NVP/NARCONDAM/2019), Andaman and Nicobar Police Department (DGP/Genl/107/20/2015/5552) and the Office of the Deputy Commissioner (South Andamans) (F.No.5-5/LS/TP/2014/7554) to conduct this study.

## DATA AVAILABILITY STATEMENT

Data will be archived in the Dryad Digital Data repository upon acceptance of this article.

## Notes

### Competing Interest Statement

The authors have declared no competing interest.

