## Supplementary Table 1 for "Frugivory and seed predation of fishtail palm (*Caryota mitis* Lour.) on the remote oceanic island of Narcondam, India"

**SUPPLEMENTARY MATERIAL**

**Table S1.** Details of the focal tree species. Stem density is for adult trees with GBH ≥ 10 cm and regeneration density is for saplings with GBH < 10 cm and height > 1 cm from 49 plots of 0.05 ha and 0.005 ha respectively.

| **Tree species** | **Family** | **Stem density ^-1^ 0.05 ha**  **(mean ± SE)** | **Regen density ^-1^ 0.005 ha**  **(mean ± SE)** | **Seed dimensions** | | | |
| --- | --- | --- | --- | --- | --- | --- | --- |
|  |  |  |  | **Seed weight (g)** | **Seed length (mm)** | **Seed width1 (mm)** | **Seed width2 (mm)** |
| *Aphanamixis polystachya* | Meliaceae | 1.87 ± 0.20 | 2.5 ± 0.50 | 0.59 | 11.32 | 9.69 | ─ |
| *Canarium euphyllum* | Burseraceae | 1.38 ± 0.26 | ─ | 1.39 | 21.61 | 12.07 | ─ |
| *Caryota mitis* | Arecaceae | 3.33 ± 0.35 | 5.73 ± 1.66 | 2.14 | 13.99 | 15.83 | ─ |
| *Chionanthus* sp. | Oleaceae | 4.27 ± 0.69 | 9.78 ± 2.21 | ─ | 31.31 | 12.87 | ─ |
| *Codiocarpus andamanicus* | Stemonuraceae | 1.88 ± 0.25 | 1 | 0.47 | 22.77 | 7.61 | 6.10 |
| *Endocomia macrocomia* | Myristicaceae | 1.48 ± 0.18 | 1 | 3.41 | 24.50 | 16.76 | ─ |
| *Planchonella longipetiolata* | Sapotaceae | 1.82 ± 0.30 | 1 | ─ | 34.01 | 15.08 | 10.74 |

The reported seed dimensions are the mean values of measurement from 5 seeds.

Seed length (in cm) measured from the base of the seed to the apex

Seed width1 (in cm) measured as longest axis perpendicular to the axis of the length

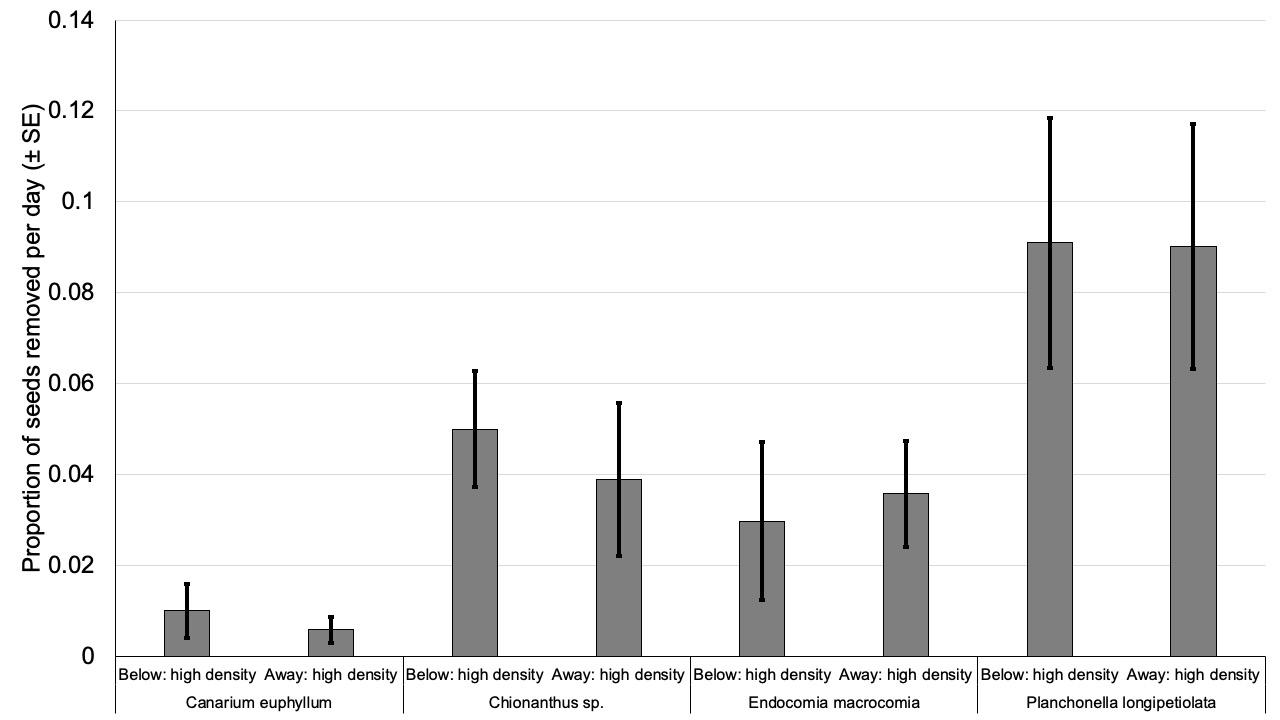

**Figure S1.** The proportion of seeds removed per day (± SE) for four species of large-seeded plants (seed width > 15 mm) on Narcondam Island. Ten seeds were laid out under the canopy of the parent tree (Below: high density) and 15 m away from the parent tree (Away: high density). The seeds were monitored for a maximum period of 33 days (mean = 15.9 days, range = 6 - 33 days) from the day of setting up the seed predation plots. The number of trees below which the seed traps were laid were three individuals for *Canarium euphyllum*, *Chionanthus* sp. and *Endocomia macrocomia* and five individuals for *Planchonella longipetiolata*.

**Table S2.** The details of the sampling effort for the seven tree species. All the tree species had one plot under the canopy and one plot away from the canopy at a distance of 20 m (10 seeds per plot). In the case of *Caryota mitis*, two additional plots were set up, one away from the canopy with low density (5 seeds per plot) and one exclosure plot under the canopy (10 seeds per plot).

| **Tree species** | **Num of trees** | **Num of plots** | **Num of seeds** | **Days monitored** | | **Num of CT** | **Days monitored** | |
| --- | --- | --- | --- | --- | --- | --- | --- | --- |
|  |  |  |  | **Min** | **Max** |  | **Min** | **Max** |
| *Aphanamixis polystachya* | 1 | 2 | 20 | ─ | 27 | 1 | ─ | 14 |
| *Canarium euphyllum* | 3 | 6 | 60 | 19 | 33 | 2 | 24 | 33 |
| *Caryota mitis* | 15 | 60 | 525 | 20 | 41 | 15 | 25 | 33 |
| *Chionanthus sp* | 3 | 6 | 60 | ─ | 12 | 2 | 12 | 12 |
| *Codiocarpus andamanicus* | 1 | 2 | 20 | ─ | 25 | 1 | ─ | 13 |
| *Endocomia macrocomia* | 3 | 6 | 60 | 12 | 14 | 2 | 12 | 12 |
| *Planchonella longipetiolata* | 5 | 10 | 100 | 6 | 26 | 1 | ─ | 13 |
| Total | 31 | 92 | 845 | ─ | ─ | 24 | ─ | ─ |

**Table S3.** Species-wise summary of proportion of seeds intact, predated and removed with respect to plot types. The three treatments are Canopy, Away high-density and Away low-density. Canopy and Away high density had 10 seeds each and Away low-density had 5 seeds. Away high-density and Away low-density plots were placed 15 m from the focal tree in opposite directions.

| **Tree species** | **Num of trees** | **Plot type** | **Prop. intact** | **Prop. predated** | **Prop. removed** |
| --- | --- | --- | --- | --- | --- |
| *Aphanamixis polystachya* | 1 | Away-high | 0.10 | 0.20 | 0.70 |
|  |  | Canopy | 0.30 | 0.00 | 0.70 |
| *Canarium euphyllum* | 3 | Away-high | 0.83 | 0.17 | 0.00 |
|  |  | Canopy | 0.73 | 0.07 | 0.20 |
| *Caryota mitis* | 15 | Away-high | 0.75 | 0.04 | 0.21 |
|  |  | Canopy | 0.81 | 0.01 | 0.17 |
|  |  | Away-low | 0.84 | 0.00 | 0.16 |
| *Chionanthus* sp. | 3 | Away-high | 0.53 | 0.47 | 0.00 |
|  |  | Canopy | 0.40 | 0.60 | 0.00 |
| *Codiocarpus andamanicus* | 1 | Away-high | 0.00 | 1.00 | 0.00 |
|  |  | Canopy | 0.00 | 0.90 | 0.10 |
| *Endocomia macrocoma* | 3 | Away-high | 0.53 | 0.47 | 0.00 |
|  |  | Canopy | 0.60 | 0.40 | 0.00 |
| *Planchonella longipetiolata* | 5 | Away-high | 0.08 | 0.92 | 0.00 |
|  |  | Canopy | 0.08 | 0.86 | 0.06 |
